# Elicitation of potent serum neutralizing antibody responses in rabbits by immunization with an HIV-1 clade C trimeric Env derived from an Indian elite neutralizer

**DOI:** 10.1101/2020.09.15.297697

**Authors:** Rajesh Kumar, Suprit Deshpande, Leigh M. Sewall, Gabriel Ozorowski, Christopher A. Cottrell, Wen-Hsin Lee, Lauren G. Holden, Sara T. Richey, Antra Singh Chandrawacar, Kanika Dhiman, Ashish, Vivek Kumar, Shubbir Ahmed, Nitin Hingankar, Naresh Kumar, Kailapuri G Murugavel, Aylur K Srikrishnan, Devin Sok, Andrew B. Ward, Jayanta Bhattacharya

## Abstract

Evaluating the structure-function relationship of viral envelope (Env) evolution and the development of broadly cross-neutralizing antibodies (bnAbs) in natural infection can inform rational immunogen design. In the present study, we examined the magnitude and specificity of autologous neutralizing antibodies induced in rabbits by a novel HIV-1 clade C Env protein (1PGE-THIVC) *vis-à-vis* those developed in an elite neutralizer from whom the *env* sequence was obtained that was used to prepare the soluble Env protein. The thermostable 1PGE-THIVC Env displayed a native like pre-fusion closed conformation in solution as determined by small angle X-ray scattering (SAXS) and negative stain electron microscopy (EM). This closed spike conformation of 1PGE-THIVC Env trimers was correlated with weak or undetectable binding of non-neutralizing monoclonal antibodies (mAbs) compared to neutralizing mAbs. Furthermore, 1PGE-THIVC SOSIP induced potent neutralizing antibodies in rabbits to autologous virus variants. The autologous neutralizing antibody specificity induced in rabbits by 1PGE-THIVC was mapped to the C3/V4 region (T362/P401) of viral Env. This observation agreed with electron microscopy polyclonal epitope mapping (EMPEM) of the Env trimer complexed with IgG Fab prepared from the immunized rabbit sera. While the specificity of antibodies elicited in rabbits associated with neutralizing autologous viruses were distinct to those developed in the elite neutralizer, EMPEM analysis demonstrated significant changes to Env conformations when incubated with polyclonal antibody sera from the elite neutralizer, suggesting these antibodies lead to the destabilization of Env trimers. Our study not only shows distinct mechanisms associated with potent neutralization of sequence matched and unmatched autologous viruses by antibodies induced in rabbits and in the elite neutralizer, but also highlights how neutralizing antibodies developed during the course of natural infection can impact viral Env conformations.

**Author Summary:** The interplay between circulating virus variants and broadly cross neutralizing polyclonal antibodies developed in a subset of elite neutralizers is widely believed to provide strategies for rational immunogen design. In the present study, we studied the structural, antigenic and immunogenic properties of a thermostable soluble trimeric protein with near native pre-fusion conformation prepared using the primary sequence of an HIV-1 clade C *env* isolated from the broadly cross neutralizing plasma of an elite neutralizer. This novel SOSIP Env trimer demonstrated comparable antigenic, structural and immunogenic properties that favoured several ongoing subunit vaccine design efforts. The novel clade C SOSIP induced polyclonal neutralizing antibody response developed in rabbits not only differed in its epitope specificity compared to that elicited in natural infection in presence of pool of viral quasispecies but also showed how they differ in their ability to influence Env structure and conformation. A better understanding of how vaccine-induced polyclonal neutralizing antibody responses compares to responses that developed in natural infection will improve our knowledge in designing better vaccine design strategies.

## Introduction

The elicitation of protective immune response by vaccination to protect against the enormous genetic diversity of HIV remains a challenge (1–5). Envelope (Env) spikes, which facilitate HIV entry and establish infection, are being considered as candidate immunogens because they mimic the trimer spike on virions (6–9). A number of recently published studies have demonstrated how structure guided stabilized Env trimers can induce potent neutralizing antibodies in different animal models (10–22). In addition, Env trimers have been used as antigen baits for the isolation bnAbs by B cell sorting and enabled structural characterization of antibody epitopes on Env (16, 23–27). Although in general, most of the trimeric Envs with closed conformation elicitated of neutralizing antibodies to tier-1 and tier 2 sequence matched autologous viruses in different animal models, their target specificities varied subtly (14, 19, 28–31). As the quality and specificity of different Env trimer-induced neutralizing antibody responses varies, presumably because of difference in the *env* sequences used to prepare the trimeric Env proteins, there is value in producing additional recombinant HIV trimers from different clades, particularly those isolated from individuals who developed broadly neutralizing antibodies. We previously reported characterized genetic and neutralization properties of *env* sequences obtained from an Indian elite neutralizer (G37080) whose plasma antibodies demonstrated >90% neutralization breadth when tested against a large heterologous Env-pseudotyped virus panel (32).

In the present study, we examined the structural, antigenic and immunogenic properties of an HIV-1 trimeric Env SOSIP protein (referred to as 1PGE-THIVC) prepared using the sequence of one of the autologous *envs* (PG80v1.eJ19) obtained from an elite neutralizer. The autologous virus is sensitive to some existing bnAbs and plasma neutralizing antibodies developed in this individual as reported earlier, but not to non-neutralizing antibodies and sCD4 (32, 33). Our overall goal in this study was to compare differences in antibody responses between immunization with recombinant 1PGE-THIVC in rabbits and those that develop during natural infection course. Three of the four rabbits immunized with the highly stable, well-ordered near native 1PGE-THIVC with closed conformation and with desirable antigenicity elicited neutralizing antibodies that demonstrated potent neutralization of tier-2 autologous virus variants, including one highly resistant *env* (PG80v2.eJ38) that was associated with escape from humoral antibody response mounted in the elite neutralizer (32, 33). Notably, neutralizing antibodies induced in rabbits by 1PGE-THIVC targeted discontinuous amino acids in the C3/T362 and V4/P401 regions on viral Env, which are distinct from those reported earlier for other Env SOSIPs. Epitopes at the C3/V4 region of HIV-1 Env targeted by SOSIP-induced neutralizing antibodies in rabbits and guinea pigs have also been previously reported (16, 34), however their antibody-specificity differed to what we have observed in this study. Moreover, we did not find (28, 29, 35) glycan holes associated with induction of SOSIP-induced neutralizing antibody response (34) in our study. The neutralizing antibody specificity developed in rabbits, however, was distinct to the kind elicited in the elite neutralizer from whom 1PGE-THIVC primary sequence was obtained.

## Results

### Characterization of an HIV-1 clade C Env trimer obtained from an Indian elite neutralizer

We previously reported an Indian elite neutralizer (G37080) infected with HIV-1 clade C whose plasma antibodies demonstrated neutralization of over 90% of a cross-clade pseudotyped virus panel (32). Additionally, we reported the degree of susceptibility of pseudotyped viruses expressing primary *envs* obtained from G37080 donor to both autologous plasma antibodies as well as different neutralizing and non-neutralizing mAbs (32, 33). One of these autologous *envs*, PG80v1.eJ19, when expressed as pseudotyped virus was susceptible to a range of neutralizing antibodies but not to sCD4 or to non-neutralizing mAbs, including V3 epitopes (3074 and 3896), b6, F105 and 17b (33). This Env was also resistant to PGT121, PGT128 and PGT135 bnAbs presumably due to natural absence of N332 at the V3 base (33). Interestingly, in contrast to all other autologous Envs, PG80v1.eJ19 was the only variant that was naturally sensitive to PGT145 (33) and hence was selected for preparation as a recombinant SOSIP trimer (1PGE-THIVC). PGT145 exclusively binds to a conformational epitope on trimeric Env, thus providing advantages in purifying pure and near native soluble Env trimers via affinity chromatography (36). Codon optimized 1PGE-THIVC SOSIP was designed, expressed in 293F or Expi293 cells and trimeric fractions were purified by PGT145 affinity column followed by size exclusion chromatography (SEC) (Figure 1A & B). The 1PGE-THIVC SOSIP assembled as >95% well-ordered trimer populations by 2D negative stain EM (Figure 1D). While 1PGE-THIVC showed a single gp140 band in SDS-PAGE (Figure 1C), the trimer converted into gp120 in the presence of DTT under reducing condition in SDS-PAGE (Figure 1E). One of the hallmarks of trimeric Envs that preferentially binds to neutralizing antibodies is that they tend to be efficiently cleaved compared to uncleaved or partially cleaved Envs, which generally expose non-neutralizing epitopes (13, 37–39). Our data indicated that the antigenicity of 1PGE-THIVC was correlated with the soluble trimers being efficiently cleaved.

**Figure 1.**
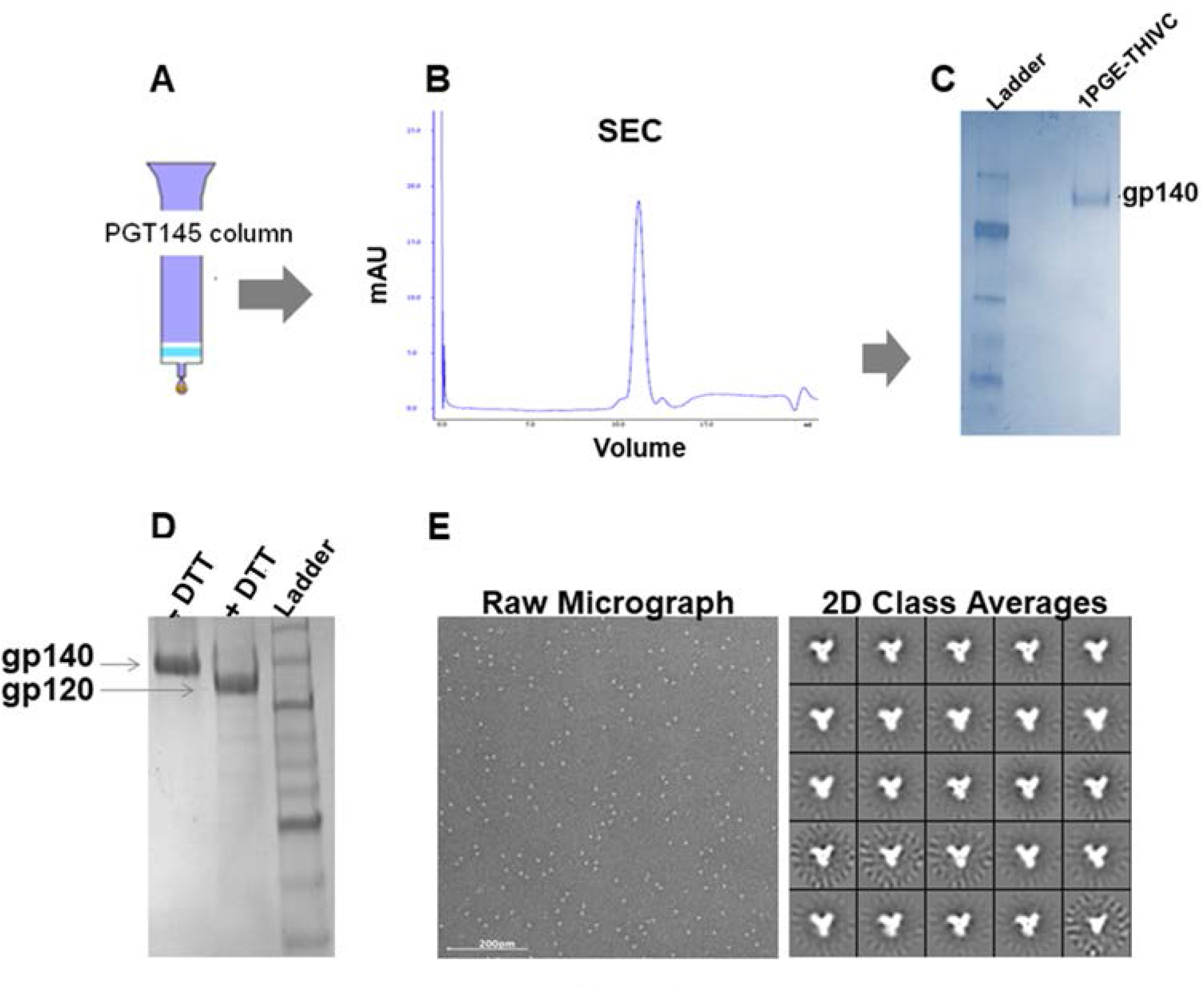
Purification of 1PGE-THIVC SOSIP. 1PGE-THIVC SOSIP was purified by PGT145 antibody affinity **(A)** and size exclusion chromatography (SEC) **(B)** followed by examination through blue native gel electrophoresis (**C)**. **D.** The SEC purified two-dimensional negative-stain EM class averages of 1PGE-THIVC SOSIP.664 trimers. **E**. Efficient cleavage of 1PGE-THIVC trimeric Env observed when run in SDS-PAGE under reducing condition.

We next examined the conformational stability of the 1PGE-THIVC Env trimer. First, we assessed the antigenicity of 1PGE-THIVC Env by measuring binding to neutralizing and non-neutralizing mAbs by ELISA (avidity). To facilitate binding with mAbs in trimeric state, D7324 epitope was introduced in the C-terminus in 1PGE-THIVC as described before (40). As shown in Figure 2A, 1PGE-THIVC preferentially bound to neutralizing mAbs over non-neutralizing mAbs. The binding affinity of 1PGE-THIVC to neutralizing and non-neutralizing mAbs with distinct specificities was next examined by BLI-Octet analysis (Figure 2B). For assessing binding affinity to bnAbs, we selected VRC01 and those that are dependent on quaternary conformation e.g., PG9, PGT145 and PGDM1400. VRC01, which targets the CD4bs, bound to 1PGE-THIVC with a very fast on-rate and very slow dissociation during wash and resulted an affinity of less than 1 nM (*KD* 1.7 nM). 1PGE-THIVC also bound strongly to PGT145 and PGDM1400 with affinities (KD) of 14 nM and 29 nM respectively (Figure 2B). Relative to PGT145 and PGDM1400, 1PGE-THIVC showed weak binding to PG9 bnAb, which also targets conformational epitopes including glycans in V1V2, with a KD of 41 nM (Figure 2B). Interestingly, the increased binding of 1PGE-THIVC to both VRC01 and CD4-Ig was found to be dependent on Asn279 (not a PNGS) on viral Env and also showed evidence in our study to significantly reduce formation of sCD4-induced higher oligomer as observed in blue native PAGE (Figure 3). We noted that the highly conserved Asn276 (glycan) is also present in 1PGE-THIVC. As expected, and in line with what we observed in binding ELISA, 1PGE-THIVC did not bind to the non-neutralizing mAb F105. Taken together, these observation results suggested that the antigenic properties of the soluble PG80v1.eJ19 SOSIP.664 Env is consistent with those expected of a well-ordered native like Env trimers as the trimer binds to neutralizing bnAbs with high affinity and has limited binding to non-neutralizing antibodies.

**Figure 2.**
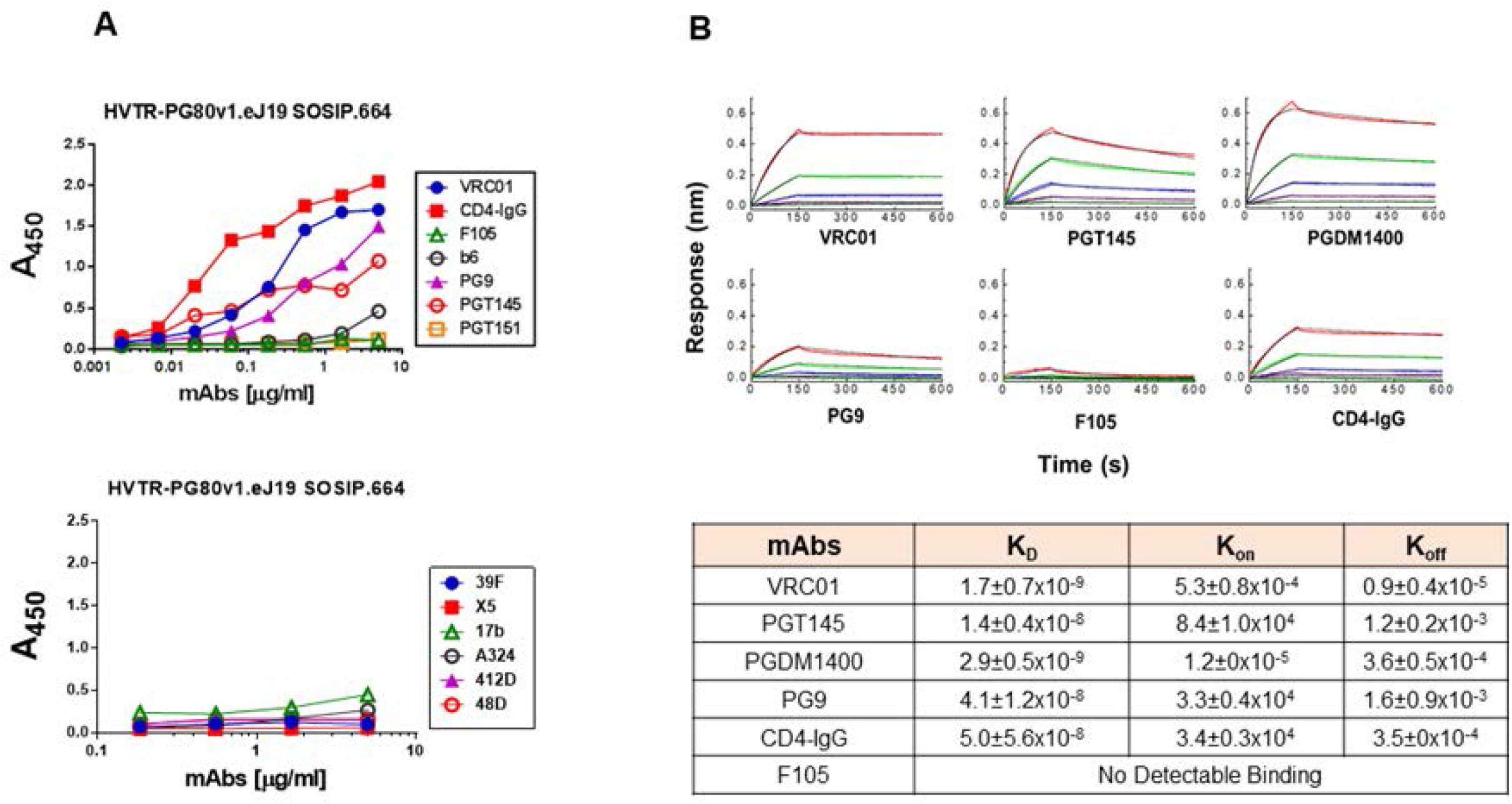
Antigenic properties of 1PGE-THIVC. **A**. Binding of 1PGE-THIVC with neutralizing and nonneutralizing mAbs by ELISA. **B.** Binding kinetics of 1PGE-THIVC to neutralizing and nonneutralizing mAbs by biolayer interferometry (BLI) kinetic analysis.

**Figure 3.**
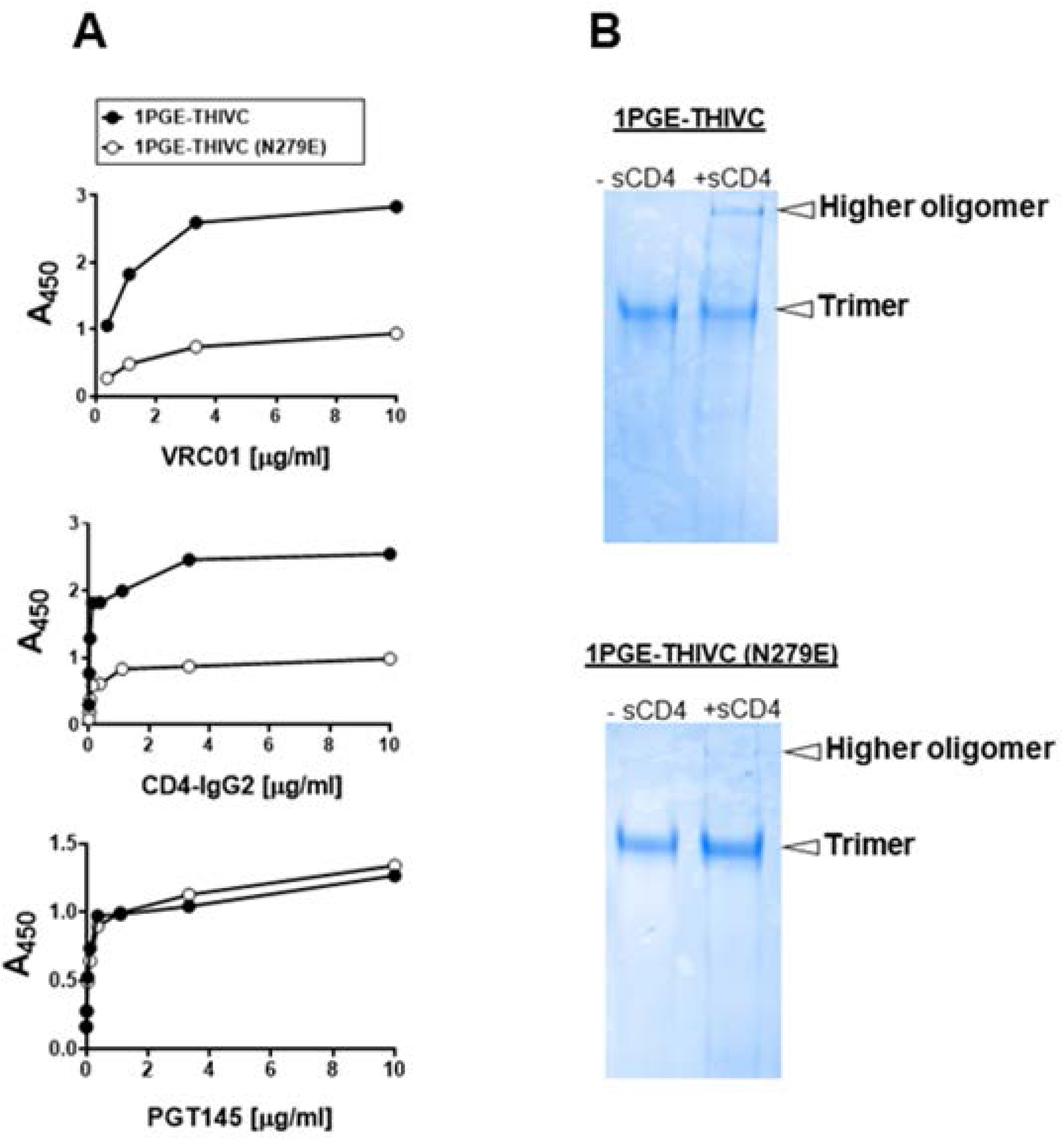
**A.** Effect of N279E on binding of 1PGE-THIVC with VRC01 and CD4-IgG2 by D-7324 capture ELISA. **B.** Blue native PAGE of SOSIP trimer (wild type and N27E version) in presence of 6-fold molar excess of sCD4.

We next examined the thermostability of the 1PGE-THIVC Env trimers by measuring the melting temperature (*Tm*) using differential scanning calorimetry (DSC). As shown in Figure 4A, the *Tm* for 1PGE-THIVC was observed to be approximately 62°C with an onset of melting at approximately 55°C. The high *Tm* obtained for the 1PGE-THIVC was comparable with the double cysteine mutant reported stable Env trimers like BG505.SOSIP.664 (13) and LT5.J4b12C SOSIP.664 (40). The soluble 1PGE-THIVC was also found to demonstrate stability at 37°C as measured by its ability to bind to different bnAbs by D7324-ELISA (Figure 4B). Overall, our data suggested that the soluble 1PGE-THIVC efficiently expresses highly stable well-ordered trimers in a closed conformation, which predominantly occludes epitopes that are targets of non-neutralizing antibodies.

**Figure 4.**
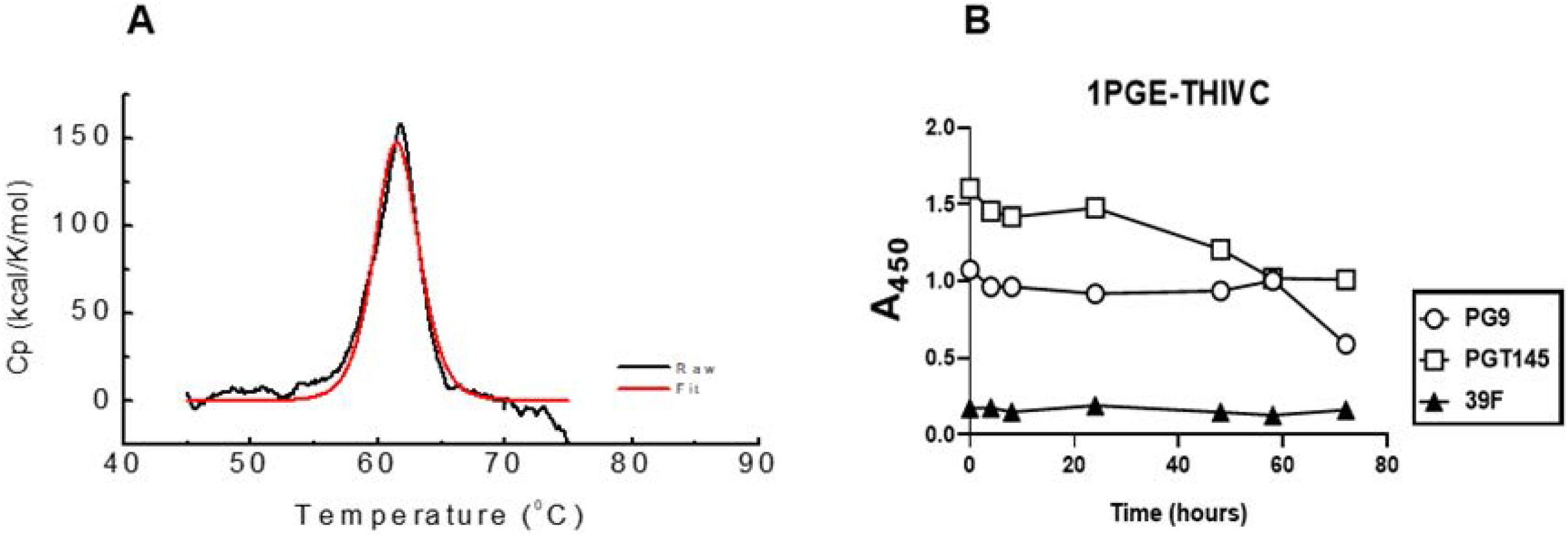
Stability of 1 PGE-THIVC SOSIP trimer. **A**. Differential scanning calorimetry (DSC) of the SOSIP trimer; *Cp*, specific heat capacity. **B.** Time course of binding of 1PGE-THIVC to bnAbs PG9, PGT145 and a non-neutralizing mAb 39F at 37°C.

### Structural properties of the trimeric 1PGE-THIVC Env as determined by Small-angle X-ray scattering (SAXS) and homology modeling

The SAXS I(q) profile of 1PGE-THIVC at concentration of 0.85 mg/ml is presented in Figure 5. The data is shown in double logarithm mode and confirms a monodisperse sample profile that lacks any aggregation or interparticulate effects. Of all the acquired data points collected (black squares), the used data points are shown in blue color. Kratky plot of the dataset show a clear peak profile supporting a globular scattering profile of the protein molecules in solution (lower inset). Distribution of interatomic vectors show that the SOSIP molecules have a maximum linear dimension (D_max_) of about 15 nm with radius of gyration (Rg) of 5.1 nm. The representative SAXS profile based on the estimation is shown as a red line in Figure 5A. The calculated SAXS data-based model of 1PGE-THIVC SOSIP molecule is shown in Figure 5B. The common envelope to all models is shown as map and the variation amongst ten models is shown as blue mesh. Normalized spatial disposition (NSD) amongst the ten models was 0.762 suggesting high similarity between the solutions. To compare this SAXS based model with previously determined structure of SOSIP, we used the primary structure of 1PGE-THIVC to generate a homology model of the protein (the closest template was PDB 6B0N) (shown as red, blue and green ribbons in Figure 5B). PDB 6B0N is crystal structure of prefusion state of HIV Env glycoprotein trimer of the clade A BG505 isolate in complex with Fabs of PGT122 and PGV19 (41). The sugar moieties of PDB 6B0N shown as magenta sticks were borrowed as such to represent glycosylation in 1PGE-THIVC. The homology model was inertially aligned over the SAXS based envelope. Three orthogonal views of the superimposition are shown in Figure 5B and 5C which provide visual confirmation that 1PGE-THIVC is also folded in P3 symmetry in same size/shape profile as previous models. Side-view shows that in the inertially aligned models, the SAXS data-based Env is not occupied in the bottom or gp41 side of the homology model (indicated by arrow). Using the SAXS and homology-based model, the different stretches of 1PGE-THIVC SOSIP as determined by SAXS and homology-based model is shown in Figure S1 (supplementary data).

**Figure 5.**
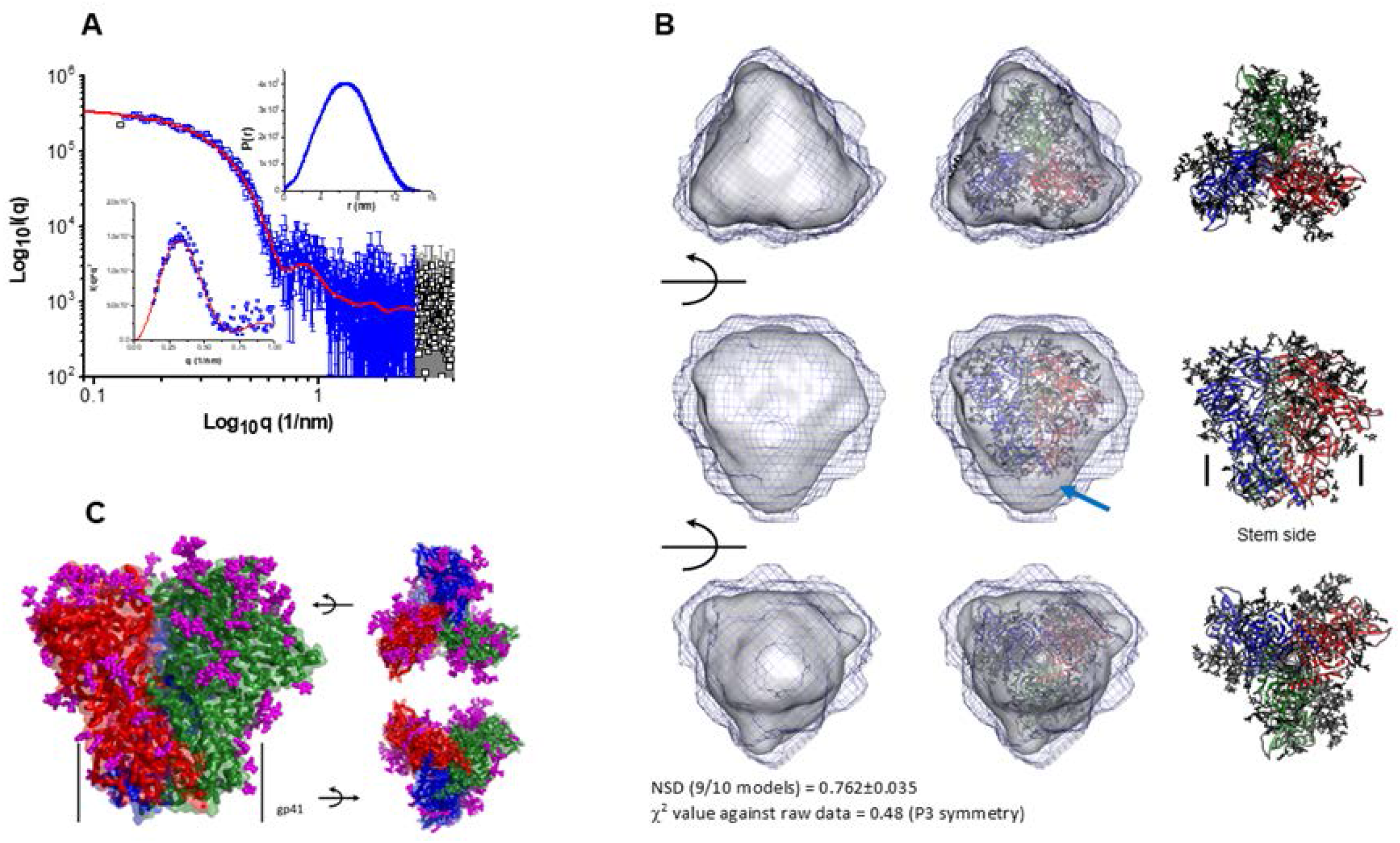
**A.** SAXS profile of the SOSIP e19 protein at about 1 mg/ml is presented in double log mode (black symbols). The blue symbols indicate the q range used to model the P(r) curve for this protein (shown as upper inset), and the red line represents the estimated SAXS profile from the P(r) curve. Lower inset shows the Kratky plot of data (blue symbols) and modelled profile (red line). **B.** (Left column) Three rotated views of the solution shape of SOSIP e19 protein restored within shape constraints present in SAXS data are shown here. Average of ten models is shown as grey map and variation in the models is shown as black mesh map. (Middle column) Automated superimposition of the residue level model of SOSIP e19 generated using primary structure of same protein and model structure of PDB ID: 6B0N with sugar moieties, and the SAXS based model have been shown. The residue level model is shown in ribbon format with three chains are shown in blue, green and red, and sugar moieties have been shown as sticks. (Right column) The residue level models superimposed in central column are shown independently. Middle panel shows where the stem side or gp41 portion is present in the model. The red arrow highlights gap in the SAXS based model vs. residue level model near the stem or gp41 region possibly due to accessible flexibility in solution. Normalized spatial disposition (NSD) amongst ten models solved using SAXS data and χ^2^ value between final averaged model vs. raw data are mentioned in bottom. **C.** Modeled structure of SOSIP e19 trimer using primary structure, SAXS data-based information and template of PDB ID: 6B0N is shown here. Three chains of gp120 are shown in red, green and blue colors. The C^α^ traces are shown as ribbons, and surfaces are shown in transparent mode. The sugar moieties representing glycosylation are shown as magenta sticks. The black arrows aid in providing the rotations done in model to present the image.

### 1PGE-THIVC Env induced potent neutralizing antibodies against sequence matched and unmatched tier-2 autologous Envs

In line with the favorable antigenic properties demonstrated 1PGE-THIVC, which is expected of well-ordered Env trimers, we next examined its ability to induce neutralizing antibodies in rabbits. Four New Zealand female white rabbits were primed and boosted with 1PGE-THIVC trimers along with Quil-A adjuvant. Two rabbits were given PBS throughout the study as control group. Quil-A was selected as an adjuvant as it was reported to stimulate the antibody-mediated immune responses against range of antigens including viruses (42), modulates antibody fine-specificity (43) and does not alter the conformation when mixed with HIV-1 Env trimers (44). The prime-boost schedule used in the process of rabbit immunization is shown in Figure 6A. Rabbits were bled before initiation of immunization (day 0) and at week 2 post priming and at weeks, 6, 8 and 12 post first boost and weeks 22, 24 and 28 following second boost with mixture of 30 μg of SOSIP protein and 40μg of Quil-A as indicated in Figure 6A. Serum samples prepared from blood samples collected at indicated intervals were heat-inactivated at 56°C for 1 hour to deactivate complement. Subsequently, the sera were evaluated for binding to 1PGE-THIVC by D7324 capture ELISA (Figure 6B) and neutralization of pseudotyped viruses expressing sequenced-matched autologous *envs* (PG80v1.eJ19 and PG80v1.eJ19 T332N) (Figure 6C & D) and two heterologous Tier 1 *envs (*SF162 and 93IN905). The peak binding of 1PGE-THIVC to serum antibodies was demonstrated by serum samples collected at week 10 and beyond from the immunized rabbits (Figure 6C). Serum samples from week 22 that demonstrated optimal binding to 1PGE-THIVC as demonstrated by D7324 ELISA was subsequently examined for its ability to neutralize sequence matched autologous Env-pseudotyped virus (PG80v1.eJ19). As shown Table 1, serum antibodies potently neutralized pseudoviruses expressing both the sequence matched (PG80v1.eJ19 and PG80v1.eJ19 T332N) and sequence unmatched (PG80v1.eJ7, PG80v1.eJ17, PG80v1.eJ158 and PG80v2.eJ38) autologous *envs.* We observed potent neutralization of PG80v2.eJ38, which was unexpected as this Env is not only resistant to autologous donor plasma antibodies (32) obtained from this elite neutralizer but also highly resistant (much like a tier-3 Env) to several bnAbs (33). Such observation was not reported earlier with any SOSIP-induced neutralizing antibodies to the best of our knowledge. Additionally, although modest neutralization of heterologous Tier 1 Env-pseudotyped viruses (SF162 and 93IN905) was observed, however with lower magnitude compared to autologous viruses (Figure 6E, F, H), which was possibly due to trimer falling apart thereby exposing the immunodominant epitopes in V3 region. Towards confirming the specificity of antibody-mediated virus neutralization, when tested, purified rabbit serum IgG was found to show neutralization of pseudoviruses expressing autologous *envs* in a dose-dependent manner (Figure 6H); thus, confirming the concordance of serum (Figure 6G) versus serum IgG mediated virus neutralization. Finally, potent autologous neutralization was correlated with efficient binding of serum IgG with 1PGE-THIVC SOSIP (Figure 6I). Taken together, our results indicated that 1PGE-THIC SOSIP trimers was able to induce antibodies in rabbits that demonstrated potent neutralization of pseudoviruses expressing sequence matched and unmatched tier-2 Envs and also demonstrated some degree of heterologous neutralization as well.

**Figure 6.**
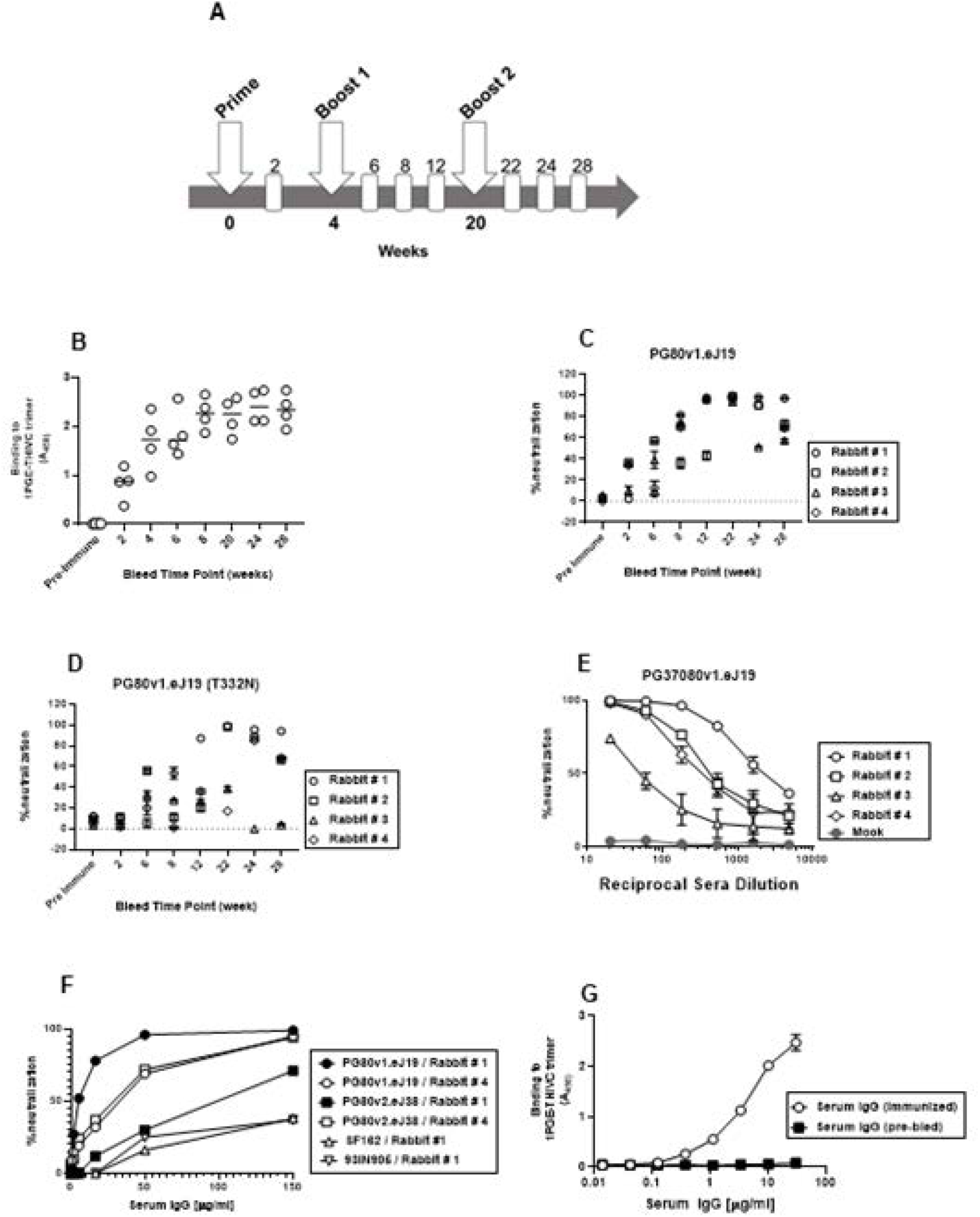
Immunogenicity of 1PGE-THIV in rabbits. **A.** Schedule of prime, boost and bleed time points. The binding **(B)** and neutralization of sequence matched (with and without N332) autologous viruses (**C and D**) by sera collected from four rabbits at 1:20 dilutions collected at different time points following prime and boost. **E.** Dose-dependent neutralization of pseudotyped virus expressing sequenced matched *env* (PG80v1.eJ19) by sera collected at week 22 following second boost with 1PGE-THIVC. **F.** Neutralization of pseudoviruses expressing sequence matched (PG80v1.eJ19) and unmatched (PG80v2.eJ38) autologous *envs* and heterologous tier1 *envs* (SF162 and 93IN905) by purified serum IgG in a dose-dependent manner; **G.** Binding of purified serum IgG (week 22; rabbit#1) to 1PGE-THIVC SOSIP trimer by D-7324 sandwich ELISA.

**Table 1.** Neutralization of sequence matched and unmatched autologous Env-pseudotyped viruses by serum antibodies obtained from 1PGE-THIVC immunized rabbits.

| **Env-pseudotyped viruses** | **Neutralization titer (ID50 values)** |  |  |  |
| --- | --- | --- | --- | --- |
|  | ***Rabbit # 1*** | ***Rabbit # 2*** | ***Rabbit # 3*** | ***Rabbit # 4*** |
| PG80v1.eJ19 WT | 2284 | 431 | 104 | 342 |
| PG80v1.eJ19 T332N | 1081 | 245 | <20 | <20 |
| PG80v1.eJ17 | 1505 | 194 | 21 | 169 |
| PG80v1.eJ158 | 1123 | 178 | 20 | 103 |
| PG80v1.eJ7 | 1037 | 141 | 20 | 161 |
| PG80v2.eJ38 | 61 | 26 | <20 | 234 |

ID50 values are reciprocal dilutions that conferred 50% neutralization of Env-pseudotyped viruses in TZM-bl cell neutralization assay.

### Mapping target epitope specificities of neutralizing antibodies elicited in rabbits induced by 1PGE-THIVC

We next examined the target specificity of the antibodies that demonstrated potent neutralization of autologous Env-pseudotyped viruses. Serum sample obtained at week-24 from rabbit #1 which demonstrated maximal and potent neutralization of autologous Envs was used to map epitope specificities using the pseudoviruses expressing wild type, chimeric and mutant autologous *env* constructs. Chimeric autologous envelope constructs were prepared between sensitive (PG80v1.eJ7 and PG80v1.eJ19) and this resistant autologous envelope (PG80v2.eJ38) as was done to assess specificities of human plasma antibodies too as shown above. As shown in Table 2, the resistant PG80v2.eJ38 Env-pseudotyped virus expressing V3/C4 sequence swapped from the sensitive PG80v1.eJ7 *env* was neutralized by serum antibodies in contrast to its wild type form, suggesting that the antibodies induced in rabbits by 1PGE-THIVC mediated potent neutralization of autologous viruses by targeting epitopes in the V3/C4 region of the viral Env. To further map its fine specificities, we prepared and tested mutant *env* constructs in the PG80v2.eJ38 backbone. We found that substitutions of amino acids asparagine (glycan) with threonine at the 362 position (N362T) in the C3 and leucine to proline at the 401 position (L401P) in the V4 regions resulted in over 8-fold increase in neutralization sensitivity (Table 2 and Figure S2) of the PG80v2.eJ38 Env-pseudotyped virus compared to its wild type form. Also, as shown in Table 2, combination of T362N and P401L substitutions resulted in over 17 and 12-fold resistance of pseudotyped viruses expressing PG80v1.eJ7 and PG80v1.eJ19 *envs* respectively having similar C3/V4 protein sequence (Figure S2) compared to their wild type forms. Interestingly, while single substitutions of T362N and P401L in PG80v1.eJ19 *env* demonstrated reduction in virus neutralization by 5.28 and 6.26-fold respectively, the combination of both demonstrated a substantial reduction in virus neutralization as described above, indicating that T362 and P401 likely comprises an epitope in the C3/V4 region targeted by neutralizing antibodies induced in rabbits. Finally, as shown in Figure 7, compared to undepleted serum, we observed a significant reduction in the neutralization of pseudoviruses expressing autologous *envs* by 1PGE-THIVC trimer-depleted serum. Taken together, our results indicate that 1PGE-THIVC SOSIP developed using a primary *env* sequence amplified from an Indian elite neutralizer induced potent autologous neutralizing antibodies in rabbits having target epitope specificities to the C3/V4 epitope on trimeric Env, which is distinct to both autologous and heterologous broadly neutralizing antibodies developed in the elite neutralizer (Table S1).

**Figure 7.**
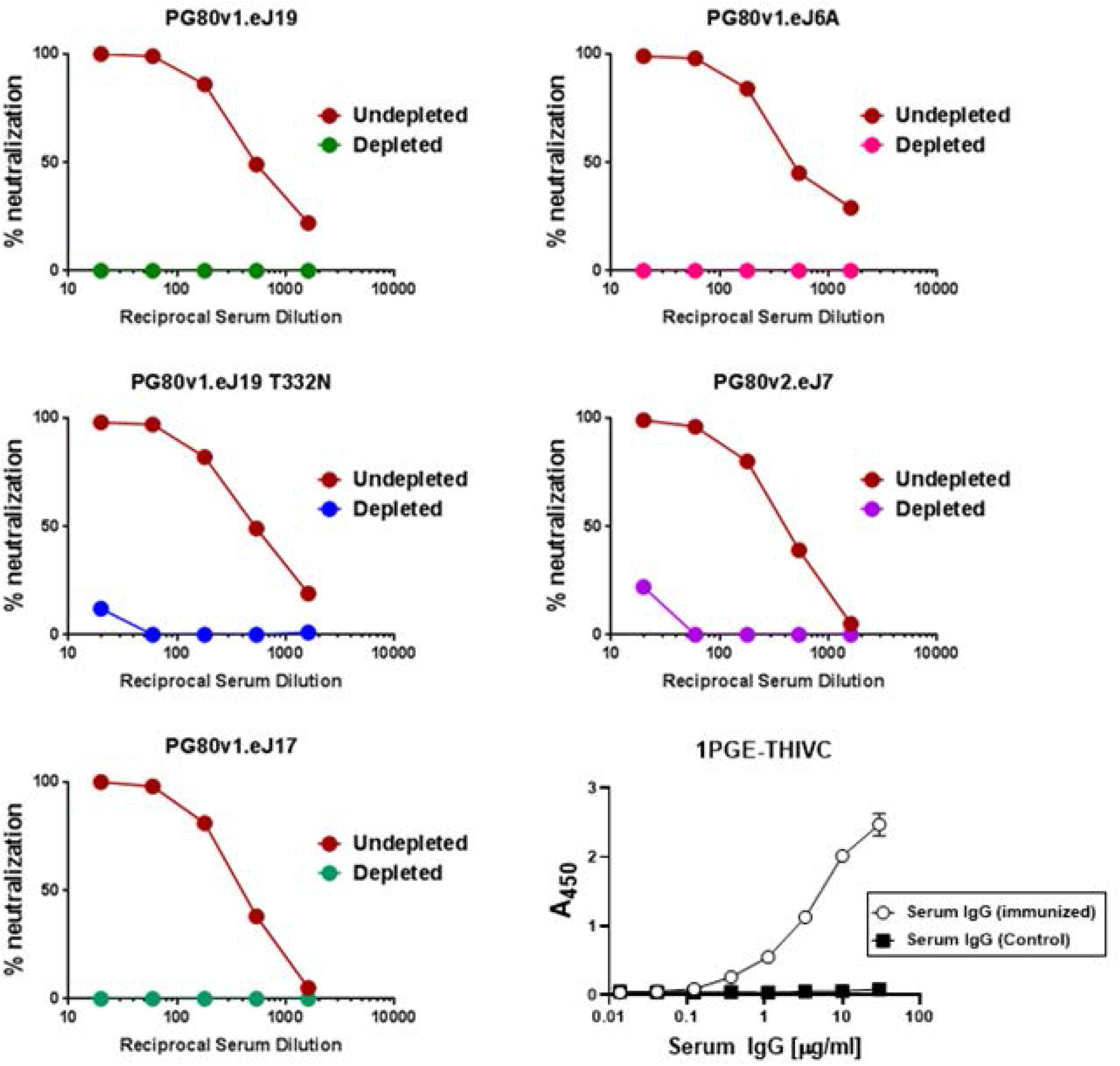
Effect of depletion of rabbit sera with 1PGE-THIVC SOSIP trimer on neutralization of pseudoviruses expressing sequence matched and unmatched autologous *envs*. Serum samples obtained at week 22 from best responder (rabbit#1) was depleted by incubating with purified magnetic beads coated with 1PGE-THVC trimeric Env. The depleted and undepleted serum was assessed for its ability to neutralize pseudoviruses expressing autologous *envs* by TZM-bl luciferase assay.

**Table 2.** Mapping fine specificity of rabbit serum antibodies associated with neutralization of autologous virus.

| Env constructs | ID<sub>50</sub> values | Fold change in neutralization of Env-pseudotyped viruses |  |
| --- | --- | --- | --- |
|  |  | Fold increase | Fold decrease |
| PG80v1.eJ7 wild type | 1798.6 | - | - |
| PG80v1.eJ19 wild type | 2117 | - | - |
| PG80v2.eJ38 wild type | 97.58 | - | - |
| PG80v2.eJ38 (V1/V2 of v1.eJ7) | <20 | No change | No change |
| PG80v2.eJ38 (V3/C3 of v1.eJ7) | <20 | No change | No change |
| PG80v2.eJ38 (V3/C4 of v1.eJ7) | 664.18 | 6.8 | - |
| PG80v2.eJ38 (C4/C5 of v1.eJ7) | 53.47 | No change | - |
| PG80v2.eJ38 (K360R) | 102.28 | 1.05 | - |
| PG80v2.eJ38 (L401P) | 418.46 | 4.3 | - |
| PG80v2.eJ38 (N362T+K360R) | 181 | 1.85 | - |
| PG80v2.eJ38 (N362T+L401P) | 789.68 | 8.1 | - |
| PG80v1.eJ7 (T362N+P401L) | 101.54 | - | 17 |
| PG80v1.eJ19 (T362N+P401L) | 171 | - | 12.4 |
| PG80v1.eJ19 (T362N) | 400.72 | - | 5.28 |
| PG80v1.eJ19 (P401L) | 337.96 | - | 6.26 |

*ID<sub>50</sub> values refer to reciprocal dilution that conferred 50% neutralization of Env-pseudotyped viruses in TM-bl cells.*

### Mapping polyclonal antibody specificities by ns-EMPEM analysis

The rabbit polyclonal serum antibody specificities developed over time following 1PGE-THIVC immunization were further examined by analyzing trimer-Fab complexes by EMPEM (45) (Figure 8). Trimer-specific responses were already observed at week 6 (2 weeks following the first boost), although against epitopes that are often seen in soluble Env immunizations and are considered non-neutralizing. These two responses are against the base of the trimer, which would not be inaccessible in a full-length native Env, and a region that coincides with the N611 glycan. Antibodies that target the N611 glycan have not been described, however recent work reveals that this site is under-occupied (i.e. it contains the correct PNGS consensus sequence, but the asparagine is glycosylated to a varying degree) in certain recombinantly expressed, engineered Env trimers, creating a neoepitope that is not believed to exist on native Env (46). These two non-neutralizing responses persist in later timepoints. By week 12 (8 weeks following the first boost) a third response is detected in the vicinity of the C3/V5 epitope. This is in agreement (and presumably the same response) as the C3/V4 epitope described above. Antibodies against this epitope have been described and tend to be potent autologous neutralizers with limited cross-reactivity. Finally, at week 22 (2 weeks following the second boost) a fourth epitope is detected by EMPEM against the region comprised of V1/V3 and/or V2 (the resolution of negative stain EM cannot discern such subtleties). This response may share some overlap with bnAbs that target the V3-glycan epitope, although the lack of heterologous neutralization by the serum implies that these antibodies rely heavily on the variable regions of Env and are more strain specific.

**Figure 8.**
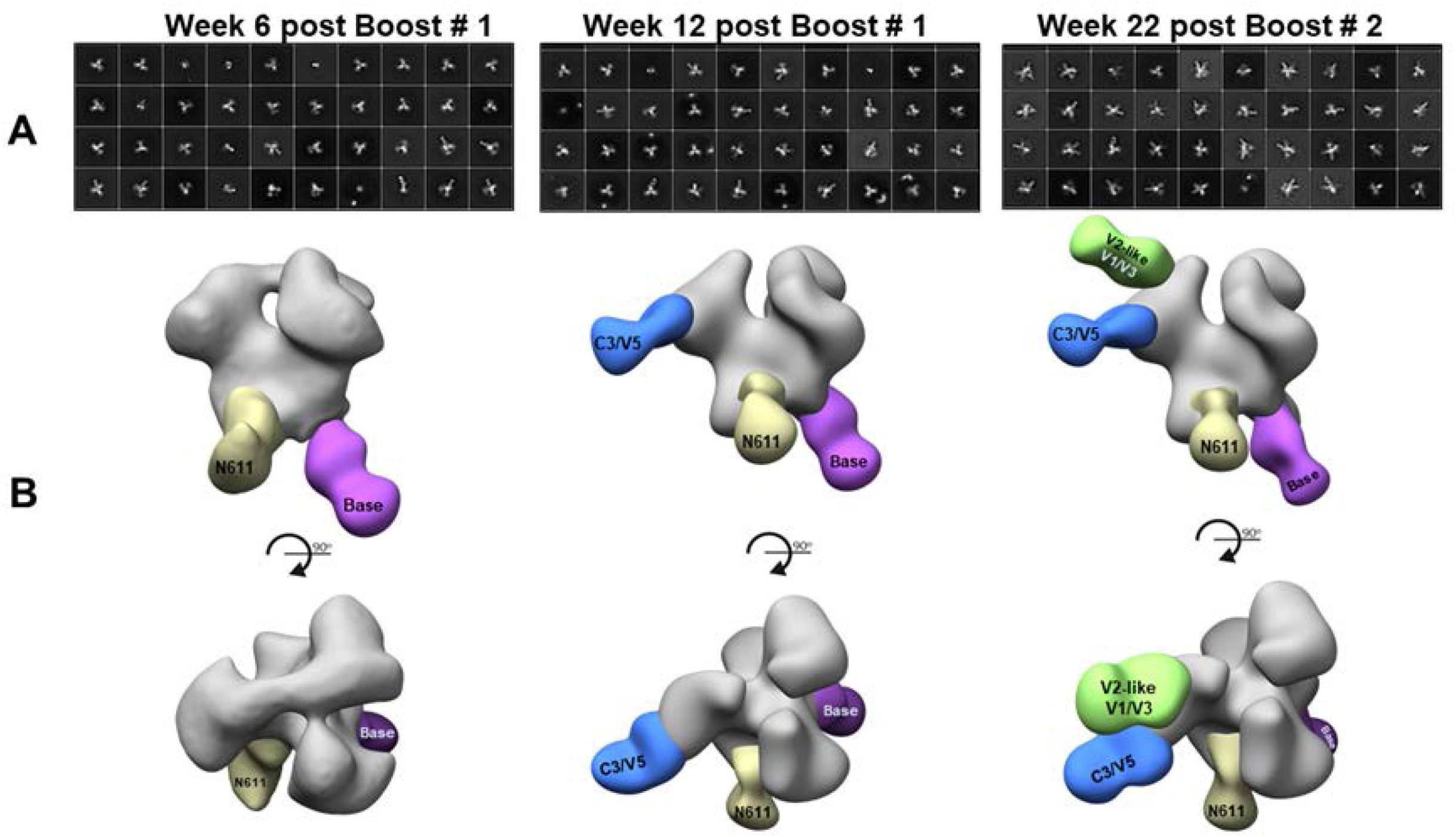
Mapping rabbit polyclonal antibody specificities by ns-EMPEM analysis. **A**. Select 2D class averages and **B.** 3D reconstructions of 1PGE-THIVC SOSIP in complex with polyclonal rabbit IgG Fab following 3 different immunization time points. For clarity, maps have been segmented into trimer (gray) and Fab (colored by epitope), and only one Fab per epitope is shown on each trimer.

### Evidence of destabilization of SOSIP trimer conformation by autologous polyclonal antibodies elicited during natural infection course in the elite neutralizer (G37080)

We next examined the impact of polyclonal plasma antibodies on the conformational state of the 1PGE-THIVC trimers. Thus, as a comparison to the EMPEM analysis we carried out with rabbit serum as described above, we also analyzed the polyclonal IgG isolated from this chronically infected human elite neutralizer that demonstrated broad cross-clade neutralization (32). Digested Fab prepared from polyclonal plasma samples were used to prepare grids with the trimer-Fab complexes as monodisperse particles. As shown in Figure 9, contrary to the rabbit immunization experiment, in which a defined set of epitopes were observed against the trimer, the EM results for the human polyclonal plasma antibodies showed a high level of heterogeneity. 2D classification revealed heterogeneity in trimer-Fab complex, in which the majority of the classes appear to be Env protomers decorated with multiple Fabs or non-native open trimers. In fact, most of the 2D class averages reveal clusters of Fabs bound to fragments of Env, with no clear indication that intact trimers are part of these complexes. In the single 2D class containing an intact trimer, no Fabs are bound (Figure 9B). These data suggest that the human donor developed antibodies that recognize multiple epitopes and were likely associated with destabilization of the engineered Env trimer ectodomain. The particles from the 2D classes do not reconstruct into an interpretable 3D density when provided a closed and ligand-free trimer as the initial model, which is likely a result of trimer dissociation (classes represent free protomers with several Fabs bound), trimer opening, or a combination of both. At least two broadly-neutralizing antibodies have been reported that induce trimer dissociation by targeting gp41, 3BC315 (derived from natural human infection) (47), and 1C2 (derived from rabbit immunization with engineered Env) (48). It has also been reported that antibodies directed against the CD4 binding site, V3 loop and the MPER can induce gp120 shedding (49). More recently, a study revealed that some V3 loop-targeting macaque antibodies can neutralize certain tier 2 viruses but require open conformations of Env with V3 exposed (50). While we are not able to assign specific epitopes from the EM data, it is clear that the responses during natural infection vary greatly comparted to the rabbit immunization study.

**Figure 9.**
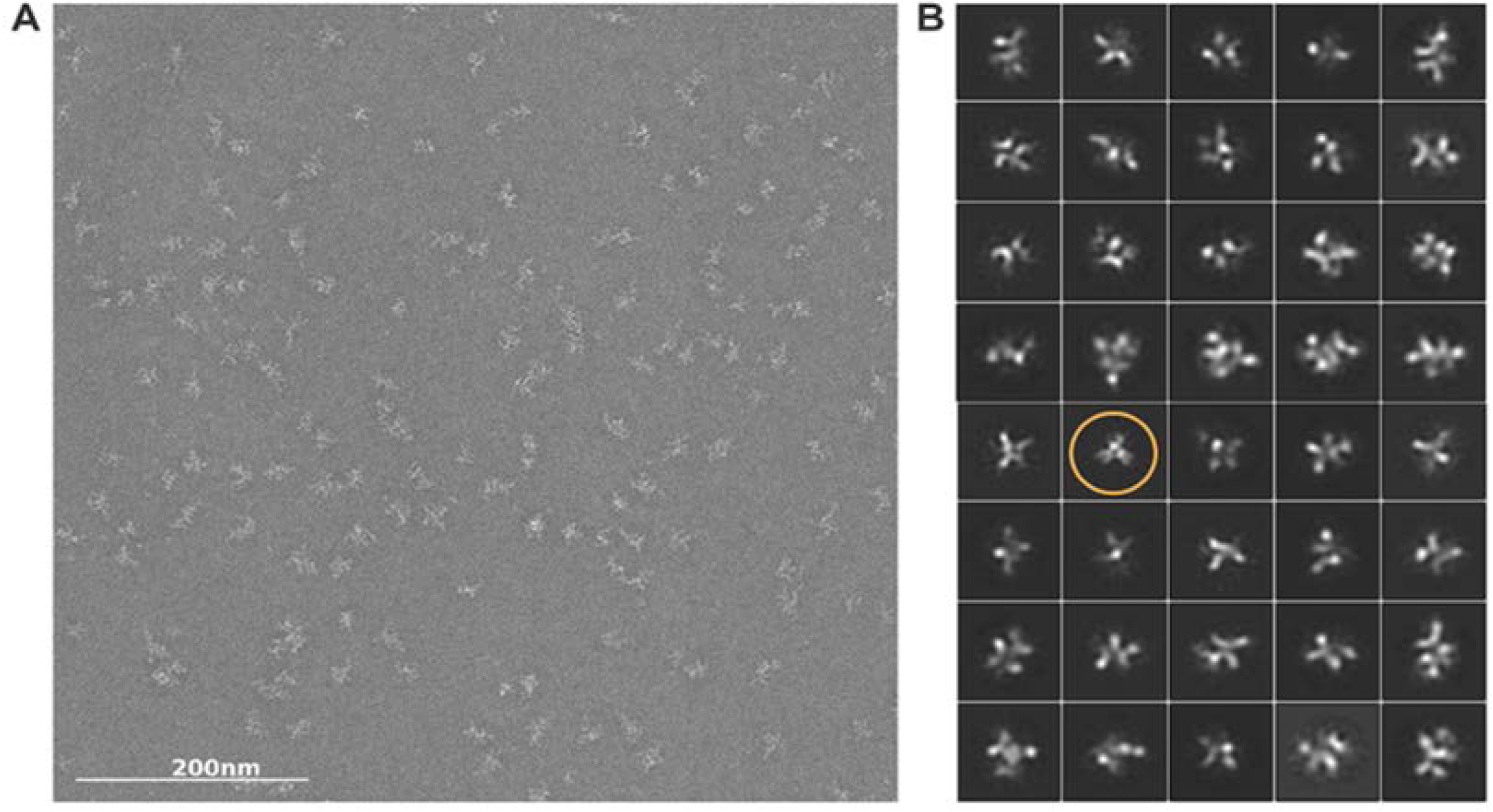
EMPEM analysis of polyclonal IgG obtained from human elite neutralizer G37080. (**A**) Representative raw micrograph and (**B**) 2D class averages reveal a high level of heterogeneity with clusters of Fabs bound to fragments of Env, with no clear indication that intact trimers are part of these complexes. A single 2D class containing an intact trimer with no Fabs bound is circled in orange. The remaining classes might represent trimer dissociation (classes represent free protomers with several Fabs bound), trimer opening, or a combination of both.

## Discussion

While near-native soluble Env trimers have been described by other studies, it remains unclear how viral envelopes obtained from individuals who developed broadly cross neutralizing antibody responses in the course of natural infection can contribute to the elicitation of bnAbs by vaccination (19). The autologous neutralizing antibody response driving virus escape is an important step towards the initiation of a cascade of viral and B cell evolutionary events resulting in the development of potent and broad neutralizing antibodies in certain individuals. Investigating the quality, magnitude and specificity of the neutralizing antibody responses induced by geographically divergent *env* sequences obtained from elite neutralizers would likely provide key strategic clues in better formulating HIV immunogens for eliciting the desired vaccine-induced protective antibody responses. Attempts have been made to examine antigenic and immunogenic properties of Env trimers prepared from *env* sequences obtained from elite neutralizers (20, 60, 61). The near native 1PGE-THIVC SOSIP soluble trimeric gp140 protein prepared using the wild type (native) *env* sequence from the Indian elite neutralizer was found not only to express very efficiently but also demonstrated the elicitation of potent autologous tier-2 neutralizing antibodies in rabbits.

The rationale for selecting the clade C PG80v1.eJ19 sequence to prepare the 1PGE-THIVC SOSIP trimer is due to its sensitivity to neutralizing and resistance to all non-neutralizing antibodies tested including those that target coreceptor binding sites, sCD4 (33) and donor serum antibodies, which indicated that the viral envelope has favorable properties that might mimic the closed pre-fusion conformation. In the present study, we sought to examine how neutralizing antibodies induced in rabbits by 1PGE-THIVC representing PG80v1.eJ19 compare in their specificity, magnitude and quality with those elicited during the course of infection in this elite neutralizer.

Before initiating the rabbit immunization, we confirmed that the thermostable 1PGE-THIVC demonstrated favorable biophysical, biochemical, antigenic and structural properties. Without any additional modification, efficient expression of trimeric Envs was observed following PGT145 affinity purification and the trimer also showed closed near native conformation by low resolution negative stain EM analysis. Moreover, an Asn279Glu substitution was found to enhance binding of the trimer to VRC01 without promoting CD4-induced conformational rearrangement that results in the open conformation of Env, which is known to expose unwanted immunodominant epitopes and induce non-neutralizing antibody response following immunization. It is intriguing that substitution of Asn279Glu, which is a part of the loop D, is very conserved (www.hiv.lanl.gov) and also forms a contact site for CD4bs directed mAbs such as HJ16 and VRC01 (62–64), demonstrated enhanced binding of 1PGE-THIVC trimer to VRC01. Removal of Asn279 position (N279) was previously shown to demonstrate detrimental effect on recognition of VRC01 class antibodies (65). The 279 position in the gp120 loop D is predominantly occupied by either Asn or Asp (www.hiv.lanl.gov) and recently was reported by LaBranche *et al.* (64) that Asn279Lys along with Gly458Tyr mutations conferred transmitted/founder virus and Env protein with enhanced susceptibility to antibodies representing germline antibodies.

We also report on how the 1PGE-THIVC folds and breathes in solution under non-frozen conditions. One of the reasons why we were keen to examine how the SOSIP trimer 1PGE-THIVC behaves in solution was to predict the conformation expected *in vivo* in circulation post rabbit immunization. The SAXS data indicated that the 1PGE-THIVC SOSIP is folded in P3 or 3-fold symmetry however retains some degree of inherent molecular mobility/disorder which possibly could result in influencing recognition by neutralizing antibodies.

The 1PGE-THIVC immunized rabbit sera neutralized pseudoviruses expressing both sequence matched and unmatched autologous *envs.* These autologous envelopes obtained from the plasma sample of the elite neutralizer at the baseline visit, although sensitive to the follow up plasma neutralizing antibodies obtained from this donor (32), were resistant to non-neutralizing mAbs including those that target the V3 immunodominant epitopes (such as 3074 and 3869) and the coreceptor binding site (e.g., 17b) (33). Interestingly, serum antibodies obtained from two immunized rabbits (rabbit # 1 and rabbit # 4) were found to show modest neutralization of the autologous PG80v2.eJ38 env that showed complete resistance to human plasma antibodies obtained in the follow up visit from the elite neutralizer (32) and to several neutralizing and non-neutralizing mAbs (33), a property that is expected of envelopes of Tier 2/3 category. Previously, we reported that PG80v2.eJ38 escaped plasma autologous plasma antibodies by mutations in V1V2 region (32); however did not distinguish the different antibody classes circulating in this elite neutralizer. In the present study, we dissected the antibody class developed in this elite neutralizer by distinguishing their specificities by depletion assays and found that the class that demonstrated exclusive autologous neutralization also had V1V2 specificity. These findings further confirmed our earlier observation of autologous virus escape by mutations in V1V2 region. In the present study, the immunized rabbit serum antibody specificity associated with neutralization of autologous viruses was mapped to C3/V4 specificity, which is distinct to the responses observed for human plasma neutralizing antibodies described above. Interestingly, although the N362 glycan residue present in the PG80v2.eJ38 *env* associated with some resistance to rabbit antibodies (and hence used for preparing chimeric env constructs for mapping antibody specificity), all were found to neutralize pseudovirus expressing PG80v2.eJ38 *env* except for the serum sample obtained from rabbit # 3. This observation is in sharp contrast to that observed with human plasma antibodies obtained from the elite neutralizer (32), which indicated that presence of proline residue at the 401 position in the C4 region possibly played a compensatory role that resulted in the neutralization of this tier 2/3 envelope. The 1PGE-THIVC induced antibodies in rabbits with C3/V4 specificity neutralized autologous virus (pseudotyped virus expressing PG80v2.eJ38) that was highly resistant to autologous human plasma antibodies as reported earlier (32), albeit with lower magnitude. Interestingly, the wild type PG80v1.eJ19 *env* lacks a glycan residue at the 332 position (N332) in the V3 base, instead contain threonine (T332). While preparing 1PGE-THIVC, we incorporated T332N, however it did not appear to induce N332 directed antibodies, neither did it improve sensitivity of the PG80v1.eJ19 T332N envelope to rabbit sera over PG80v1.eJ19 T332 envelope. We are unable to comprehend the basis for resistance of PG80v1.eJ19 T332N envelopes to serum samples obtained at week 22 from rabbit # 2 and 3 while the same sera neutralized pseudovirus expressing the PG80v1.eJ19 T332 *env*.

Despite production and *in vitro* characterization of well-ordered native like Env trimers, one of the concerns that remains post immunization is the stability and the ability of the Env trimers to retain its integrity *in vivo* as subtle distortion in conformation can lead to the induction of non-neutralizing antibodies via off-target binding. Loss in virus neutralization by trimer-depleted serum antibodies indicated that 1PGE-THIVC induced autologous neutralizing antibodies conformational epitopes. Indeed, serum antibodies obtained at week 22 from the rabbit that demonstrated the most potent neutralization also neutralized heterologous envelopes including those are categorized as Tier 1A. The magnitude of heterologous neutralization also differed; for example, SF162 and MW965 which are of Tier 1A category (66) showed maximal neutralization sensitivity compared to 93IN905 envelope which is of Tier 1B category (14). Intriguingly, this was not noticed with serum samples obtained from early time points post immunization. It is also important to note that the tier-2 clade A envelope, Q259.d2.17 was neutralized by the same serum antibodies at the same time. Therefore, it is unknown whether the neutralization of Tier 1 envelopes was exclusively mediated by non-neutralizing (such as those are directed to immunodominant V3 epitopes).

To further understand the quality of polyclonal antibodies developed in immunized rabbits, the polyclonal antibody imaging approach by negative stain EM analysis was carried out with serum samples collected at weeks 6 and 12 following first protein boost and at week 22 following second protein boost. The common feature of all these three serum samples is the presence of two non-neutralizing responses; gp41 base and a N611 glycan hole. The former is seen in majority of the soluble trimers that have been examined as this high peptide content base is highly immunogenic. The sub-occupancy of a glycan at position N611 has been demonstrated in BG505 SOSIP using mass spectrometry and cryo-EM (46). The N611 PNGS is highly conserved across clades (www.hiv.lanl.gov) and both PG80v1eJ19 and 1PGE-THIVC contain the consensus sequence for glycosylation at this position N611. While this is believed to be true for soluble SOSIP Env proteins, it is necessarily not true for primary viruses (67). Interestingly, in contrast to BG505 SOSIP, which demonstrated elicitation of strong 241/289 “glycan hole” directed antibody responses in rabbits (35), our data suggest that 1PGE-THIVC Env while this clade C *env* sequence naturally blocks those glycan epitopes. Additionally, rabbit sera also neutralized the pseudotyped virus expressing both parental PG80v1.eJ19 *env* having T332 and also its T332N version (Table 1), further indicating that neutralizing antibodies induced in rabbits also did not target N332 glycan epitope too. The serum samples collected at weeks 12 and 22 demonstrated a C3/V5 directed antibody response, similar to previously published reports (34). This epitope is predicted to be dependent on glycan repositioning in the V5 loop, and since the V5 glycosylation site(s) vary across genotypes, cross-reactivity is likely to be hampered as described very recently (68). Finally, the serum sample collected at week 22 following second SOSIP boost demonstrated a fourth antibody response by nsEMPEM analysis that indicated “V1/V3” and “V2-like” antibody responses, which at low resolution (2D classification) remains difficult to distinguish. However, it should be noted that neither responses appeared the same as that of canonical bnAb “V1/V2-apex” or “V3-glycan supersite.” We believe therefore that these antibodies will potentially have high levels of strain specificity as they interact with the V1, V2 and/or V3 variable regions. Of note 1PGE-THIVC SOSIP trimers appear “more open” at the apex in the 3D reconstructions compared to BG505 SOSIP. What this means immunologically is hard to say since there do not appear to be responses to newly exposed epitopes, nor are epitopes of the four distinct antibodies responses measured against 1PGE-THIVC SOSIP known to induce opening. The V3 signal observed in nsEMPEM could partially explain neutralization of tier 1 envelopes, however, neither neutralization of tier 1 envelopes was observed with serum samples up to week 12 post SOSIP boost nor any elicitation of V3 directed antibody observed by polyclonal EMPEM analysis. Interestingly, in contrast to the specificities of neutralizing antibodies induced in rabbits as observed by EMPEM analysis, broadly cross neutralizing polyclonal antibodies developed in the human elite neutralizer drastically altered the trimer conformation of the sequence matched antigen that was used for rabbit immunization. While such observation has not been documented previously in the context of polyclonal antibody mapping, our observation indicates that a population of antibodies in the polyclonal mixture, possibly correlated to the broad neutralization developed in this elite neutralizer, are capable of destabilizing trimers, exposing additional epitopes that are normally buried surfaces (for example though gp120-induced shedding in natural infection) as shown in other unrelated studies (47, 49). This clearly suggests that the neutralizing antibody responses induced in rabbits by 1PGE-THIVC SOSIP trimer are different than those elicited during the course of natural infection in this individual. Although not known definitively, such phenomenon may possibly be linked with development of virus-antibody co-evolution leading towards achieving neutralization breadth. While in contrast to the rabbit antibodies, we could not infer human polyclonal antibody specificity by EMPEM, our data highlights how antibodies developed in natural infection that are tied neutralization breadth can significantly impact upon Env conformation. It would be important to further investigate the mechanism underlying this phenomenon by examining virus and antibody interaction by following individuals infected with HIV overtime, particularly those who go on to develop cross neutralizing antibodies in the course of natural infection. In summary, it is important to study different well-ordered native SOSIPs in different animal models, which would better inform what structural, antigenic and immunogenic properties can better guide and select Env SOSIPs that would likely to help achieve neutralization breadth to genetically diversified HIV. Moreover, since subtype C accounts for nearly half of the global HIV infection (www.hiv.lanl.gov), it is important to select and study region-specific HIV-1 Envs (e.g., clade C) and which are associated with mounting neutralizing antibodies *in vivo* in the natural infection course. Such exercise will likely be able to inform the rational design and development of immunogens that can least be able to largely dissect the region-specific diversity of circulating HIV-1 subtypes such as clade C.

## Materials and Methods

### Ethics Statement

The blood samples were collected under the IAVI Protocol G study from slow-progressing ART naive HIV-1-positive donors from Nellore District of the state of Andhra Pradesh, southern India, by trained clinicians at the YRG Care Hospital following approval and clearance from the Institutional Review Board (IRB) and the Ethics Committee. The plasma samples collected were shipped to the Translational Health Science and Technology Institute, for research purpose only. The rabbit immunization work was outsourced to a contract research organization that obtained necessary approvals from animal ethics committee prior to conducting immunization and collecting blood samples from vaccinated rabbits.

### Preparation of 1PGE-THIVC SOSIP trimer

Codon optimized HIV-1 Indian clade C (1PGE-THIVC) gp140 SOSIP trimeric protein was prepared based on a Tier-2 HIV-1 clade C primary *env* sequence (PG80v1.eJ19), obtained from an Indian elite neutralizer (32) essentially as described by Sanders *et al.* (13). 1PGE-THIVC gene was codon-optimized by GeneArt (Thermo Fisher Inc.) and cloned into pcDNA3.1(+) with following modifications to the wild-type Env sequence: A501C, T605C, I559C (for trimer stabilization), and gp120 - gp41 cleavage motif REKR changed to RRRRRR. The D7324 epitope sequence (GSAPTKAKRRVVQREKR) was added after residue 664 in gp41 ectodomain (ECTO) and preceding the stop codon to facilitate examining Env SOSIP binding by ELISA following published protocol (40, 51). 1PGE-THIVC was expressed by transient transfection of 293F or Expi293 cells, and the trimeric protein fraction was purified from culture supernatants first by using PGT145 mAb affinity column (36). Bound proteins were eluted with 3 M MgCl_2_, dialyzed with PBS pH (7.4), and subsequently concentrated using Amicon ultracentrifuge filters (Millipore) with a 100-kDa cutoff to 0.5-1 ml. The PGT145 affinity purified SOSIP protein was further subjected to size exclusion chromatography (SEC) using a HiLoad Superdex 200 16/60 column (GE Healthcare Inc.). The SEC purified proteins were snap frozen in liquid nitrogen and stored at −80°C until further use. Purified trimeric SOSIP proteins were analyzed in a gradient 4-15% BN-PAGE (Mini-PROTEAN TGXTM, Bio-Rad). The degree of 1PGE-THIVC cleavage was examined by incubating the SOSIP protein with 0.1 M dithiothreitol (DTT) and analyzed by SDS-PAGE under reducing conditions as described earlier (40).

### Trimer ELISA

Binding of SOSIP trimers to different mAbs by D7324 sandwich ELISA was carried out as described previously (40). Briefly, high-binding microtiter plates (Nunc, Inc.) were first coated with D7324 antibody (Aalto Bio Reagents, Dublin, Ireland) at 10 μg/ml (100 μl/well) in coating buffer (150 mM Na2CO3, 350 mM NaHCO3, 30 mM NaN3, pH 9.6) at 4°C overnight. Microtiter plates were washed three times using phosphate buffered saline (PBS) with 0.05% Tween-20 and subsequently blocked with 220 μl of 5% (w/v) nonfat milk in PBS and incubated at 37°C for 1 hour. Purified 1PGE-THIVC-D7324 trimers were added at 500 ng/ml in PBS (100 μl/well) for 2-3 hours. Unbound trimers were removed by washing three times with PBS. PBS containing 3% (w/v) skimmed milk (250 μl/well) was subsequently added to block nonspecific protein-binding sites. The ELISA binding reactions were initiated by incubation of SOSIP protein to varying concentrations of mAbs for 1 hour at 37°C. After three washes with PBS, 100 μl of anti-human HRP (Jackson ImmunoResearch Laboratories Inc.) diluted at 1:2000 was added and incubated at room temperature for 50 min. The plates were further washed four times with PBS containing Triton X-100 (0.05% v/v) and color developed by addition of 100 μl of tetramethylbenzidine (TMB) substrate. Absorbance was measured at 450 nm in an ELISA reader (BioTek Inc.).

### Biolayer Interferometry

For binding kinetics anti-human Fc sensors (Octet, ForteBio Inc.) were used to capture the mAbs, whereas SOSIP trimer was used as analyte in varying concentrations (ranging from 210 to 2.6 nM) in the HEPES buffer background supplemented with 0.02% Tween 20 and 0.1% BSA (pH 7.2). The binding of antigen (SOSIP) and antibody (mAbs) were carried out in room temperature by incubation of SOSIP-bound biosensors in wells containing mAbs (10 μg/ml) for 120 s with agitation at 1000 rpm. Binding association was recorded for 150 s followed by dissociation for 450 s. Data were analyzed using the ForteBio Data Analysis software, 9.0 (Forte-Bio Inc) and using a 1:1 binding model to fit the association and dissociation curves. A global fit was performed using all curves in which the concentration of SOSIP yielded a change in binding of at least 0.1 nm and a measurable dissociation.

### Differential Scanning Calorimetry (DSC)

SOSIP protein in PBS (pH 7.2) diluted to 0.1-0.2 mg/ml was loaded onto a Micro-Cal VP-Capillary DSC instrument (Malvern, Inc.) and subjected to a 20 - 90 °C ramp at 60°/ h. Origin 7.0 software was used to subtract baseline measurements and to fit the melting curves using a non-two-state model. Reported T_m_ values are for the tallest peak of each sample.

### Negative stain EM

1PGE-THIVC SOSIP trimers were diluted to 0.01-0.03 mg/ml, applied to a carbon coated Cu400 grid, and stained with 2% (w/v) uranyl formate as described previously (19). Data were collected on an FEI Tecnai Spirit T12 transmission electron microscope operating at 120 keV and equipped with a Tietz TVIPS CMOS camera. A magnification of 52,000x was used, resulting in a physical pixel size at the specimen plane of 2.05 Å. Data processing and analysis methods have been reported elsewhere (19). Two-dimensional classifications were performed using MSA/MRA (52).

### Small Angle X-Ray Scattering (SAXS)

All SAXS experiments described here have been performed on SAXSpace instrument (Anton Paar GmbH, Austria). The instrument had a sealed tube X-ray source, a line collimated X-ray beam and a 1D CMOS Mythen detector (Dectris, Switzerland). The wavelength of X-rays was 0.154 nm and the sample to detector distance was about 317.6 mm. SAXS data was acquired on three samples of SOSIP at concentrations of 0.72, 0.85 and 1.1 mg/ml. For each concentration, the sample was exposed for 60 minutes (2 frames of 30 minutes each) at 10°C in a thermostated quartz capillary with diameter of 1 mm. The scattering data captured at detection was re-calibrated for the beam position using SAXStreat software. The SAXSquant software was then used to subtract buffer contribution, set the usable q-range, and desmear the data using the beam profile. The SAXS data was further analyzed using the programs available in the ATSAS 2.7 suite of programs (53). The radius of gyration (Rg) was calculated on the basis of automated Guinier approximation using the PRIMUSQT integrated suite of programs (54). The Porod Exponent, x was estimated by plotting I(q)*q^x^ vs q^x^ till the profile resembled hyperbolic profile. The molecular mass of the scattering particles/protein molecules was calculated using the DATMOW program. Same suite was used to compute the distance distribution function in auto-mode using the program GNOM which performs an Indirect Fourier transformation on the SAXS intensity profile (55). *Ab initio* models were generated using first computing ten independent models using DAMMIF program, superimposed and averaged using SUPCOMB and DAMAVER programs, and then the averaged structure was refined using DAMMIN program. Calculations of DAMMIF were done considering no (P1) or P3 symmetry. The SAXS based model of SOSIP e19 was compared with its PDB 6B0N based homology model by inertial axes alignment of two models using SUPCOMB program. For structural visualization and figure generation, open source Pymol and UCSF Chimera programs were used.

### Preparation of Env-pseudotyped viruses

HIV-1 Env-pseudotyped viruses were prepared as described previously (32). Briefly, 293T cells were co-transfected with envelope-expressing plasmid and an *env*-deleted HIV-1 backbone plasmid (pSG3ΔEnv) using a FuGENE6 transfection kit (Promega Inc.). Cell supernatants containing pseudotyped viruses were harvested 48 h post transfection and used for infection in TZM-bl cells using DEAE-dextran (25 μg/ml) in 96-well microtiter plates. The virus infectivity titers were determined by measuring the luciferase activity using Britelite luciferase substrate (PerkinElmer Inc.) in a luminometer (Victor X2, PerkinElmer Inc.).

### Site-directed mutagenesis

Point mutations by site-directed mutagenesis were introduced in *env* constructs using the QuikChange II kit (Agilent Technologies Inc.) following the manufacturer’s protocol. Introduction of desired substitutions was confirmed by sequencing as described previously (32).

### Neutralization assay

Neutralization assays were carried out using TZM-bl reporter cells as described before (56). Briefly, Env-pseudotyped viruses were incubated with varying dilutions of antibodies (mAbs, serum and plasma) for 1 h at 37°C in a CO2 incubator under humidified condition. TZM-bl cells (1 X 10^4^) were added into the mixture virus-antibody mixture containing 25 μg/ml DEAE-dextran (Sigma). The plates were further incubated for 48 h and the extent of virus neutralization was assessed by measuring relative luminescence units in a luminometer (Victor X2, PerkinElmer Life Sciences).

### Rabbit immunization

New Zealand white female rabbits were immunized with 30 μg of 1PE-THIVC SOSIP formulated with 40 μg Quil-A adjuvant (Invivogen Inc.) at weeks 0, 4 and 20. Four rabbits were taken in antigen immunized group and three rabbits were taken in placebo group. In placebo group, animals received PBS (pH 7.0). Pre-bled sera were obtained from all the rabbits prior to immunizations and bleed samples were collected from each animal at different time point as mentioned in Figure 6. Serum samples obtained post boost 2 were assessed for their extent to neutralize autologous and heterologous Env-pseudotyped viruses. The rabbit immunization was outsourced to a contract research organization (CRO) at Bengaluru, Karnataka, India.

### Depletion of plasma and serum antibodies by monomeric gp120 and trimeric SOSIP proteins

Purified soluble monomeric (92BR020 gp120) and trimeric 1PGE-THIVC (SOSIP gp140) proteins, in were used for the depletion of human plasma and rabbit serum neutralizing antibodies as described earlier (32). Briefly, purified gp120 and SOSIP proteins were covalently coupled to 30 mg of tosylactivated MyOne Dynabeads (Life Technologies Inc.) in coupling buffer [0.1 M NaBO_4_,1M(NH_4_)_2_SO_4_; pH 9.4] overnight at 37°C for 16 to 24 h according to the manufacturer’s protocol. Env proteins bound to magnetic beads were separated from unbound proteins using a DynaMag 15 magnet (Life Technologies, Inc.). Env protein bound beads were further incubated with blocking buffer (PBS [pH 7.4], 0.1% bovine serum albumin [BSA; Sigma], and 0.05% Tween 20) at 37°C to block the unbound sites and the antigenic integrity of Env proteins were assessed by examining their ability to bind to different mAbs by flow cytometry (FACSCanto; Becton and Dickinson, Inc.). For depletion studies, plasma and serum samples were diluted to 1:50 in Dulbecco’s modified Eagle’s medium (DMEM) containing 10% fetal bovine serum (FBS), and 500μl of diluted plasma or serum were incubated with 20μl of Env protein coupled magnetic beads at room temperature for 45 min. Unbound plasma and serum antibodies were separated from bound antibody fraction using a DynaMag 15 magnet as described above. This step was repeated 4 to 5 times for gp120 and 1-12 times for SOSIP protein (1PGE-THIVC) towards facilitating efficient depletion. As a negative control, G37080 plasma antibodies were depleted with uncoated beads in parallel. In addition to ELISA, the percent depletion of G37080 plasma. The degree of depletion of the polyclonal serum and plasma antibodies were assessed by ELISA and TZM-bl neutralization assay as described previously (32).

### Polyclonal Fab preparation

Serum immunoglobulin G (IgG) was purified with a mixture of protein A/G affinity column. Purified IgG was digested for 6 hours at 37° C using 4% (w/w) liquid papain (Thermo Fischer Scientific) and digestion buffer (10 mM L-cysteine, 10X EDTA, pH 8). The digestion solution was collected, and Fab fragments were purified from undigested IgG and Fc-fragments using SEC (Superdex 200 Increase; GE Healthcare). Final Fab yields were ~0.75-1.5 mg. Complexes were assembled with 10-15 *μ* g of 1PGE-THIVC (SOSIP gp140) trimer and ~1 mg of purified polyclonal Fab, at room temperature for 18 hours. They were then purified using SEC (Superose 6 Increase; GE Healthcare) with TBS as a running buffer and concentrated with 10 kDa cutoff Amicon ultrafiltration units. Samples were diluted in TBS to ~30 *μ* g/ml and immediately deposited onto carbon-coated 400-mesh Cu grids (glow-discharged at 15 mA for 25 s), where they were then stained with 2% (w/v) uranyl formate for 30 s. For each sample, 116,958 to 250,000 individual particle images were collected and were subsequently submitted to 2D and 3D classification using Appion (57) and Relion 3.0 (58) data processing packages. Figures were generated using UCSF Chimera (59) by aligning representative 3D reconstructions for a specific time point and animal to each other and segmenting the maps into Fab and trimer segments. For clarity, figures only display one Fab density per epitope and a single trimer density.

## DATA AND SOFTWARE AVAILABILITY

EM volumes have been deposited to the Electron Microscopy Data Bank under accession codes EMD-22496, and EMD-22498 through EMD-22505 (inclusive) (https://www.emdataresource.org/).

## Acknowledgements

We thank the Protocol G study participant (G37080) registered with YRG Care, Chennai, all of the research staff members at the Protocol G clinical center at YRG Care, Chennai, and all of the IAVI Protocol G team members. We sincerely thank all our laboratory members for providing valuable input in preparing the manuscript. The 92BR020 wild type and mutant gp120 constructs as well as several bnAbs tested in this study were provided by the IAVI Neutralizing Antibody Center, the Scripps Research Institute, La Jolla, California, USA. We also thank Prof David Montefiori for kindly providing us with the following *env* clones: Q23.17, CH038 and Q259.D22.2 used in this study. The following reagent was obtained through the NIH AIDS Reagent Program, Division of AIDS, NIAID, NIH, from John C. Kappes and Xiaoyun Wu: pSG3 env. IAVI’s work was made possible by generous support from many donors, including the Bill & Melinda Gates Foundation, the Ministry of Foreign Affairs of Denmark, Irish Aid, the Ministry of Finance of Japan, the Ministry of Foreign Affairs of the Netherlands, the Norwegian Agency for Development Cooperation (NORAD), the United Kingdom Department for International Development (DFID), and the United States Agency for International Development (USAID). The full list of IAVI donors is available at www.iavi.org. The contents are the responsibility of the International AIDS Vaccine Initiative and do not necessarily reflect the views of USAID or the United States Government. We thank Prof Gagandeep Kang, Dr Rajat Goyal and Prof Sudhanshu Vrati, for support.

## Author contribution

JB conceptualized the study; JB, RK, SD, GO, A, ABW designed the study; RK, VK designed the codon optimized SOSIP and examined biophysical and biochemical properties of the Env trimer and examined antigenic and immunogenic properties and thermostability; RK performed plasma depletion assays; RK, SD carried out neutralization assays; SD, NK prepared *env* mutant constructs and mapped neutralizing antibody specificities of rabbit and human serum samples; ASC, KD, A performed SAXS analysis of Env trimer; RK, NH prepared bulk volume of 1PGE-THIVC Env and purified serum IgG for EM structural studies at TSRI; LS, GO, CAC, WL, LGH, SR, ABW carried out EMPEM analysis; SA assisted in BLI-Octet analysis; KGM, AKS identified and recruited the human elite neutralizer under IAVI Protocol G study; KGM prepared and provided the donor plasma and serum samples used in this study in addition to providing related clinical data; DS provided reagents for antibody mapping studies and helped in data analysis. JB wrote the manuscript with help from all the authors.

## Supporting documents

**Table S1.** Mapping specificities of the autologous and heterologous plasma neutralizing antibodies obtained from the Indian elite neutralizer.

**Figure S1.** Different epitopes and loops in the primary structure of 1PGE-THIVC SOSIP trimer are highlighted in these images. In parentheses, highlighted residues and their respective colors are mentioned. The black arrows aid in providing the rotations done in model to present the image.

**Figure S2.** Alignment of C3/V4 amino acid sequences of the autologous *envs* obtained from G37080 donor. Amino acid numbering is made based on HXbc2 sequence. Amino acid residues in C3 and V4 that form key epitopes targeted by neutralizing antibodies induced in rabbits are highlighted. The glycan residue at the 362 position is underscored.

